# The Neurocognitive Architecture of Individual Differences in Math Anxiety in Typical Children

**DOI:** 10.1101/160234

**Authors:** Charlotte Emily Hartwright, Chung Yen Looi, Francesco Sella, Alberto Inuggi, Flávia Heloísa Santos, Carmen González-Salinas, Jose M. García Santos, Roi Cohen Kadosh, Luis J Fuentes

**Affiliations:** Aston Brain Centre, School of Life and Health Sciences, Aston University, Birmingham, UK; Department of Experimental Psychology, University of Oxford, Oxford, UK; School of Experimental Psychology, University of Bristol, Bristol, UK; Istituto Italiano de Tecnologia, Genova, Italia; Departamento de Psicología Básica y Metodología, Facultad de Psicología, Universidad de Murcia, Murcia, Spain; Departamento de Psicología Evolutiva y de la Educación, Facultad de Psicología, Universidad de Murcia, Murcia, Spain; Servicio de Radiología, Hospital Morales Meseguer, Murcia, Spain

## Abstract

Math Anxiety (MA) is characterized by a negative emotional response when facing math-related situations. MA is distinct from general anxiety and can emerge during primary education. Prior studies typically comprise adults and comparisons between high-versus low-MA, where neuroimaging work has focused on differences in network activation between groups when completing numerical tasks. The present study used voxel-based morphometry to identify the structural brain correlates of MA in a sample of 79 healthy children aged 7-12 years. Given that MA is thought to develop in later primary education, the study focused on the level of MA, rather than categorically defining it as present or absent. Using a battery of cognitive- and numerical-function tasks, this study identified that increased MA was associated with reduced attention, working memory and math achievement. Increased MA was associated with reduced grey matter in the left anterior intraparietal sulcus. This region was also associated with attention, suggesting that baseline differences in morphology may underpin attentional differences. Future studies should clarify whether poorer attentional capacity due to reduced grey matter density results in the later emergence of MA. Further, our data highlight the role of working memory in propagating reduced math achievement in children with higher MA.

## Introduction

Math anxiety (MA) is characterised by negative emotional response such as fear and tension when facing math-related situations, which cannot be reduced to either general anxiety or test anxiety^1^. It disrupts mathematical performance irrespective of gender^2^, and can emerge in the primary school years^3,7^. Depending on the extent of MA, the negative impact of MA could have far-reaching consequences beyond academic achievements^4^. Given that a significant variation in the level of MA is contributed by genetic factors^5^, understanding the neurocognitive basis of individual differences in MA may shed light on its causal pathway.

To date, there are a limited number of neuroimaging studies, which are mainly based on adults and comparisons of neural response between groups of high and low levels of MA. In adults, high-compared with low-level MA has been shown to be associated with increased activity in bilateral posterior insula, brain areas linked with threat and pain processing, when anticipating mathematical tasks^6^. High-compared to low-level MA has also been associated with stronger deactivation within the default mode network during tasks that require additional inhibitory functions, possibly reflecting depletion of working memory resources^7^. Further, increased activity in frontoparietal regions in high-level MA adults when anticipating mathematical tasks has been shown to correspond with reduced performance deficits, suggesting the role of cognitive control in MA^8^. In children^9^, high-level MA has been associated with hyperactivity in the right amygdala when solving mathematical problems, and increased connectivity between this and the ventromedial prefrontal cortex, areas implicated in the processing and regulation of negative emotions. Compared with low-level MA, high-level MA has been linked with decreased activity in brain areas involved in working memory and attention, including the dorsolateral prefrontal cortex, and reduced activity in posterior parietal areas, critical to numerical processing.

A recent review^10^ highlighted that there are specific aspects of numerical and executive function that might explain varying degrees of MA: lower working memory capacity, reduced attentional control, lower inhibitory control and a deficit in low-level numerical representations. Furthermore, increasing levels of MA may negatively affect mathematical achievement, via disruption of core executive functions and/or deficit in low-level numerical representations. Using data collected from a larger study, we sought to test the strength of association made by those predictions in that review, to better understand the neural and cognitive factors that are associated with the degree of anxiety towards mathematics. We aimed to identify the structural brain correlates of the *level of MA* in a typical school population, and to determine how brain structure mediates the relationship between those cognitive functions that are most strongly predictive of the level of MA. Much of the prior literature has consisted of between-group comparisons and, whilst these have provided important insights on the plausible neural mechanisms of MA, the neural and cognitive architecture that contributes to individual differences in MA, and its association with mathematical achievement, remains unclear.

The present study comprises brain structure and MA data from 79 healthy, Spanish children. The Behavioural Rating Inventory of Executive Function (BRIEF)^11,12^ was used to profile each child's behavior in specific domains of executive function. The BRIEF indices that were of primary interest for the present study were: inhibitory control and impulsivity (INHBIT), ability to switch and alternate attention (SHIFT) and on-line, representational memory (WORKING MEMORY). The BRIEF is regularly used in clinical and education settings, where higher scores indicate greater difficulty, and therefore lower capacity, in each domain. Numerical representation skills were assessed using a number line task, consisting of positioning numbers on an analogue scale (PN)^13^, mathematical achievement was determined using the Woodcock Johnson III Achievement (WJ)^14^ and the level of MA was established via the Math Anxiety Scale (Math-AS)^15^. Voxel-based morphometry (VBM) was used to identify brain-structure correlates of MA.

## Method

### Participants

Participants were recruited through two state primary schools in Murcia, Spain, as part of a wider study^20–22^. The primary sample comprised 137 Spanish children, aged 7-12 years (2^nd^ – 6^th^ grade). Consent was obtained from parents prior to acquiring any data, and reobtained immediately prior to data acquisition. T1-weighted structural MRI data were acquired from an initial sample comprising 110 children whose parents gave previous informed consent. Two were not included in the current study as they were reported to be bilingual, which may affect the measurement of math anxiety and numerical achievement. A further 7 were excluded due to learning disability. Following exclusion due to unsatisfactory image quality resulting from movement- or other imaging artefacts (*n*=21) or neuro-incidental findings (*n*=1), the final sample comprised 79 children (age *M* = 115.20, *SD* = 14.13; males = 50.6 %; right-handed = 88.6%). The study was approved by the University of Murcia Ethics Committee, and it was conducted in accordance with the approved guidelines and the Declaration of Helsinki.

### Materials

Measures that were modeled in the current study include math anxiety, mathematical abilities and working memory.

### Math Anxiety

MA was assessed using the Math-AS (known also as the EAM, Escala de Ansiedade a Matemática ^15^). It consists of 25 items that describe situations that are commonly experienced by elementary and high school students during their math lessons. This scale measures the variations in the degrees of math anxiety, from absence to extreme math anxiety. This task was translated from Portuguese into European Spanish by a speaker fluent in both languages (FHS) to enable cross-cultural adaptation. Children indicated the intensity of their response to each item on a five-point Likert scale by crossing out one of the following: (1) None (2) Low (3) Moderate (4) High and (5) Very High. The score was the sum of all points from the 25 questions. This measure was shown to have accurate validity and reliability when used in the children population ^23^.

## Numerical Cognition

### Math Achievement

Children’s math abilities were assessed using the Spanish version of the Woodcock-Johnson III (WJ-III) Achievement (ACH) battery ^14^, which has been validated for the use of children aged 6-13 years in Spain ^24^. It comprises 4 subtests: calculation, math fluency, quantitative concepts and applied problems (see *Supplementary Information: Method).* The raw scores of each subtest were transformed into W scores ^25,26^ following the Rasch’s measurement model ^27,28^. We used the composite score of all 4 subtests in our analyses.

### Numerical Representation

Numerical representation was assessed using a number line task, consisting of positioning numbers on an analogue scale (PN)^13^. The participants were required to map numbers on a vertical line that was marked with “0” at the bottom and “100”at the top. In half of the trials, the line was further marked with 4 horizontal lines at different locations to assist children with number mapping. Children were required to indicate the position of an Arabic numeral, orally or visually presented by the experimenter by pointing to a specific location on the line. There were 12 trials in this task.

### Executive Function

We used parents’ rating of children’s online memory (WORKING MEMORY), attention (SHIFT) and inhibitory control (INHIBIT) using the Spanish version of the Behavioural Rating Inventory of Executive Function, BRIEF™ ^11,12^, which assesses the executive functioning of children between 5-18 years old. It contains 86 items in eight non-overlapping clinical scales and two validity scales.

## Procedure

### Cognitive and Achievement Testing

Children’s performance on a range of behavioural, cognitive and achievement tasks was assessed prior to collecting the structural MRI scans. Testing was conducted during the Autumn term by six trained assistants with children in groups of two.

### Analysis of Demographic, Cognitive and Achievement Data

Several of the measures resulted in positively skewed data. Parametric statistics were combined with permutation testing as this approach, in contrast with non-parametric analyses, has been shown to maximally reduce type I and type II errors^29^. Confidence intervals (CIs) were estimated using the bias-corrected and accelerated (BCa) percentile bootstrap method (10,000 samples).

The cognitive and achievement data were analysed using SPSS, version 22. The mediation analysis was conducted using the Process macro for SPSS (v2.16.3), available from http://www.processmacro.org/index.html following a published analysis pipeline^30^. Ten-thousand bootstrap resamples were used to generate bias-corrected, 95% confidence intervals.

### Neuroimaging Acquisition

The participants were fitted with ear plugs and soft foam padding used to minimize head movement during the scan. Participants were asked to remain as still as possible for the duration of the scan, and a parent sat beside their child throughout. A T1-weighted image was acquired for each participant using a 1.5T GE HDX scanner with an 8-channel, phased array, transmit-receive head coil. A 3D FSPGR BRAVO sequence was used to achieve whole brain coverage, composed of 142 axially oriented slices with a reconstructed voxel size of 1×1×1mm^3^, where TR = 12.4 ms, TE = 5.2 ms, flip angle = 12°.

### Neuroimaging Analysis using Voxel-Based Morphometry

The MRI data were analyzed using the FMRIB Software Library (FSL, version 6.0.0; http://fsl.fmrib.ox.ac.uk/fsl/fslwiki/). Non-brain tissue was removed from the structural images and an initial weak bias field correction applied using FSL’s anatomy pipeline (http://fsl.fmrib.ox.ac.uk/fsl/fslwiki/fsl_anat). These brain extracted, bias corrected images were then fed into the second and subsequent stages of FSL-VBM^31^ (http://fsl.fmrib.ox.ac.uk/fsl/fslwiki/FSLVBM), an optimised VBM protocol ^32^. The images were grey matter-segmented and registered to the MNI-152 standard space using nonlinear registration^33^. The resulting images were averaged and flipped along the x-axis to create a left-right symmetric, study-specific grey matter template. All native grey matter images were then non-linearly registered to this study-specific template and “modulated” to correct for local expansion (or contraction) due to the non-linear component of the spatial transformation. The modulated grey matter images were then smoothed with an isotropic Gaussian kernel with a sigma of 3 mm, Full-Width-Half-Maximum (FWHM) ∼7 mm. Finally, voxelwise general linear modelling was applied using Randomise^34^ (http://fsl.fmrib.ox.ac.uk/fsl/fslwiki/Randomise), which permits permutation-based nonparametric testing, correcting for multiple comparisons across space. Here, 10,000 permutations of the data were generated to test against the null. Threshold-free cluster enhancement (TFCE)^35^ was used to identify cluster-like structures, taking family-wise error rate (FWE) corrected *p*-values <.05. To avoid any labelling bias, probabilistic anatomical descriptors were determined using FSL Atlas Query (http://fsl.fmrib.ox.ac.uk/fsl/fslwiki/Atlasquery). Anatomical labels were output for voxels that had survived multiple-comparison correction. Cluster peak information was extracted using FSL’s Cluster tool. See *Supplementary Information: Method* for a detailed description of the GLM analyses.

## Results

### Math Anxiety and its Association with Demographic, Cognitive and Numerical Factors

Table 1 outlines the sample’s demographic, numerical and cognitive characteristics (see also *Supplementary Information: Table S1* for a detailed breakdown by grade). Typically, the level of MA within the sample was low. An independent-samples t-test determined that there was no difference in the mean level of MA between the sexes (t(65.569) = −1.063, *p* =.292, 95% CI −13.21, 3.569). Age was positively associated with the level of MA (*r*(79) =.237, *p* =.035, CI.031,.433) (see *Supplementary Information: Fig. S1*).

To assess the relationship between MA, numerical- and cognitive-function we conducted a series of partial correlations. Each of the three BRIEF indices of interest (INHIBIT, SHIFT, WORKING MEMORY), plus the PN and WJ were correlated with the Math-AS score (controlling for age and biological sex). SHIFT, WORKING MEMORY and WJ were statistically significant after applying a Bonferroni correction (Table 2).

**Table 1.** Sample Characteristics

| **Domain** | **Scale/Index** | ***n*** | ***mean*** | ***SD*** | **min** | **max** |
| --- | --- | --- | --- | --- | --- | --- |
| **Demographic** | Age (months) | - | 115.20 | 14.13 | 95.00 | 145.00 |
|  | School grade | - | 4 | - | 2 | 6 |
|  | Sex (M/F) | 40 / 39 | - | - | - | - |
|  | Handedness (L/R) | 9 / 70 | - | - | - | - |
| **Numerical Cognition** | Math Anxiety Scale | 79 | 43.52 | 19.35 | 25 | 101 |
|  | Woodcock-Johnson III Achievement | 79 | 128.15 | 29.64 | 65 | 208 |
|  | Number line task | 73 | 7.51 | 2.18 | 2.00 | 10.50 |
| **Executive Function (BRIEF)** | Initiate | 71 | 12.92 | 2.99 | 8 | 22 |
|  | Working Memory* | 71 | 17.49 | 4.62 | 10 | 27 |
|  | Plan | 71 | 19.87 | 5.43 | 12 | 33 |
|  | Organization | 71 | 10.75 | 3.28 | 6 | 18 |
|  | Monitor | 71 | 13.63 | 3.20 | 8 | 22 |
|  | Inhibit* | 71 | 15.34 | 3.53 | 10 | 27 |
|  | Shift* | 71 | 12.69 | 3.05 | 8 | 21 |
|  | Emotional Control | 71 | 16.49 | 4.21 | 10 | 28 |
|  | Behavioral Regulation Index | 71 | 43.96 | 9.41 | 18 | 69 |
|  | Metacognition Index | 71 | 74.69 | 16.70 | 48 | 118 |
|  | Global Executive Composite | 71 | 119.21 | 24.32 | 79 | 184 |

*Note.* \*Indicates BRIEF indices of primary interest due to prior published associations with Math Anxiety

**Table 2.**
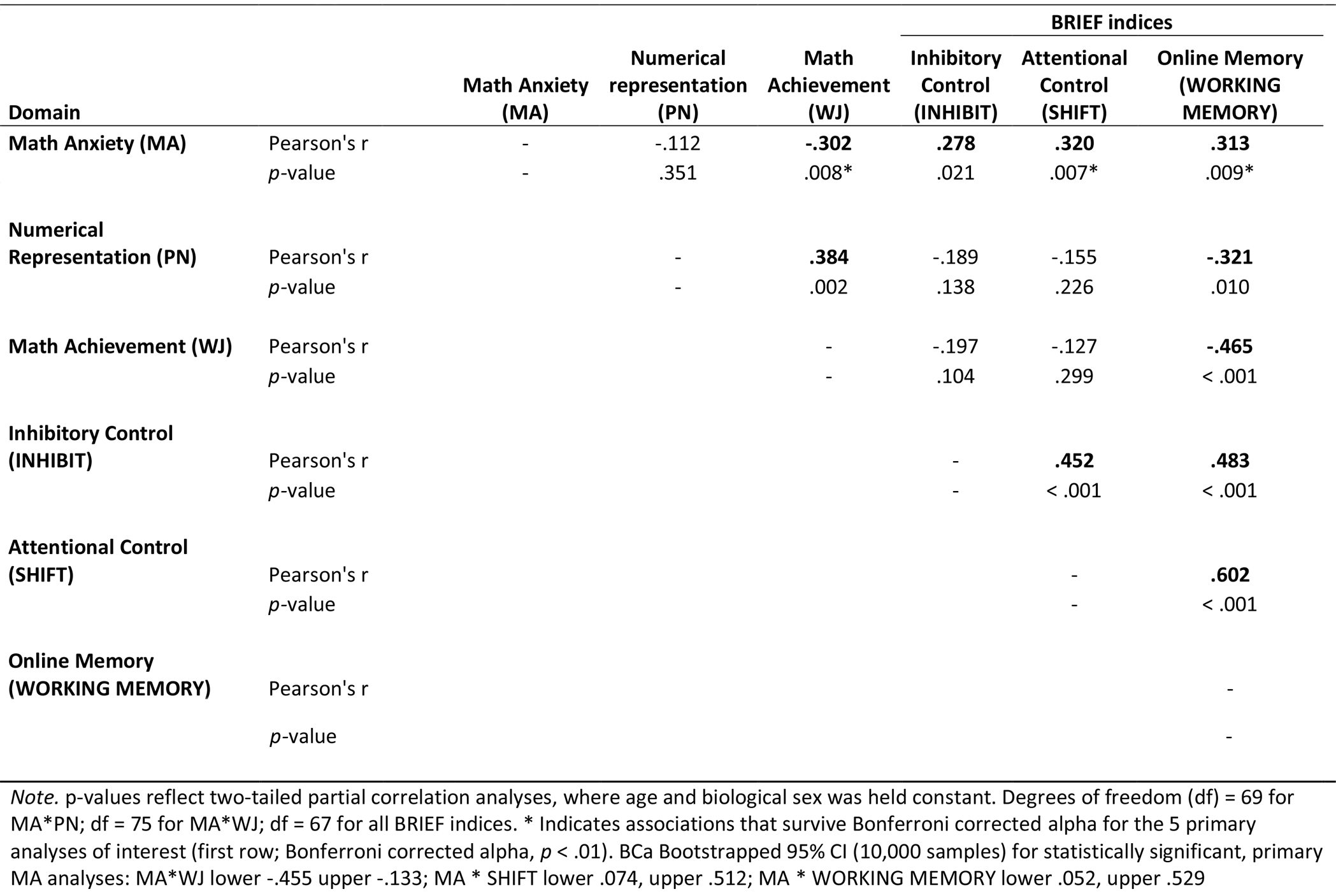
Association between Math Anxiety, Numerical and Cognitive Indices

### Math Anxiety and its Association with the Brain

VBM was used to identify the structural correlates of MA. A general linear model (GLM) comprising Math-AS score and the nuisance variables, age, biological sex, recruitment source and handedness was created. Contrasts for positive and negative associations between grey matter volume (GMV) and MA were computed, where corrections were applied for multiple comparisons across the brain, and adjustment to correct for running two contrasts using a Bonferroni correction. A whole brain analysis did not identify any regions that were positively associated with MA. Four clusters demonstrated a negative association between GMV and MA, which encompassed only cortical grey matter (Table 3 and Fig. 1). The largest cluster spanned both hemispheres across occipital and parietal cortices, including a section running anterior-posterior along the left anterior intra-parietal sulcus (IPS), in areas P1 and P3, as defined by the Juelich Histological Atlas. A second, smaller cluster was identified in the right visual cortices, encompassing extrastriate areas and additionally encompassing the left anterior IPS. Lastly, a small, left lateralized cluster was identified in the inferior parietal lobule. The Bonferroni correction had split this latter cluster, resulting in a further cluster in the lateral occipital cortex. As this comprised a single voxel (MNI: −18, −88, 42), this voxel was excluded from subsequent analyses.

**Figure 1.**
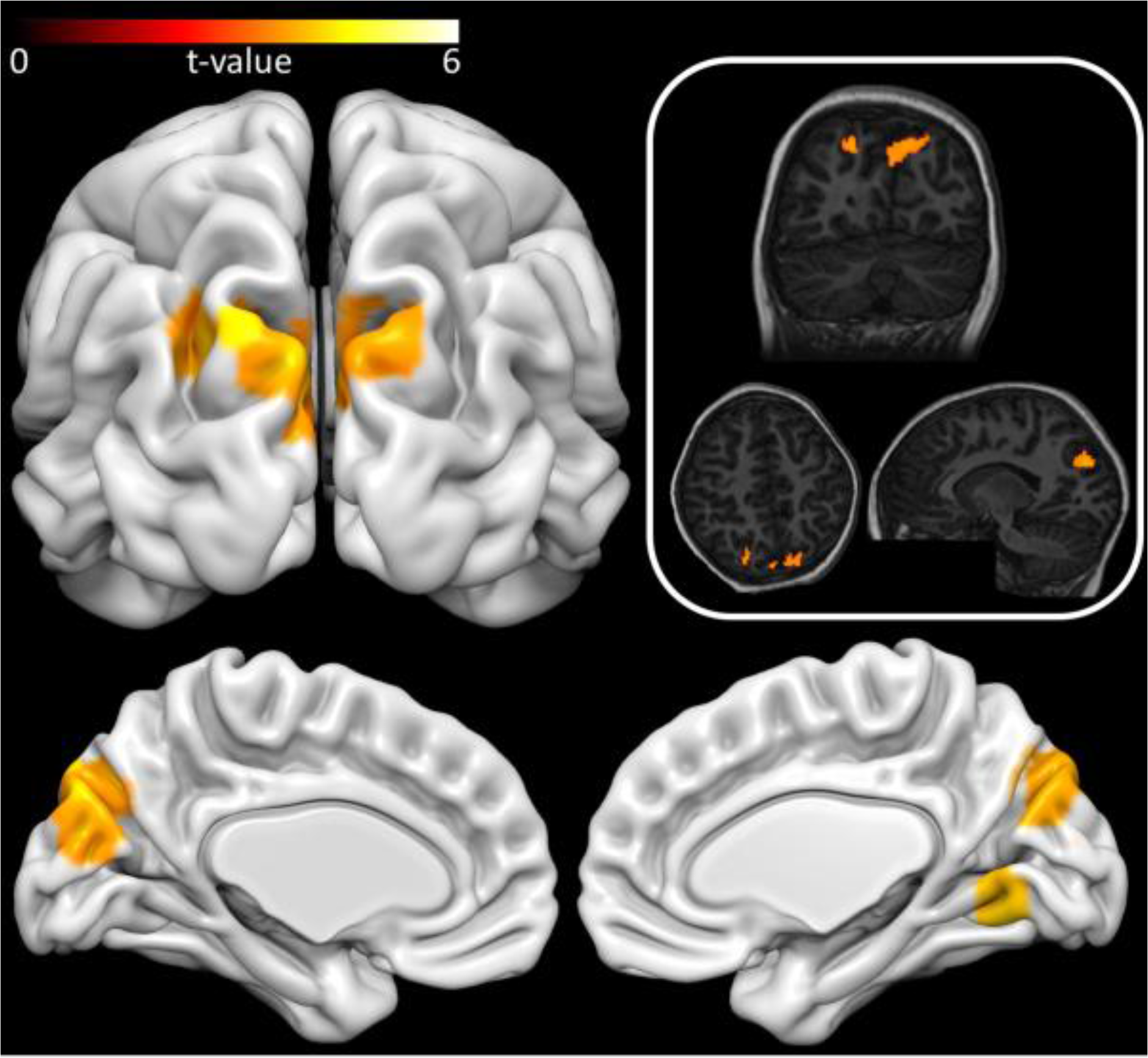
Grey matter correlates of Math Anxiety. Surface rendered image reflects a t-statistic cluster map rendered onto a template brain. All coloured areas reflect those grey matter voxels that were significantly negatively associated with MA after multiple comparison corrections as outlined in the method. The top right panel illustrates the results transformed and rendered onto a single participant’s annoymised T1-structural image. All images are presented in neurological convention, where the left of the image reflects the left of the brain. Surface rendering created using Surf Ice (v 10.11.16) [Computer software], available www.nitrc.org/projects/surfice/ Individual subject rendering created using Mango (v 4.0), [Computer software], available from http://ric.uthscsa.edu/mango/mango.html

**Table 3.** Probabilistic Labels for Brain Regions where Grey Matter Volume is Negatively Associated with Math Anxiety

| HO Anatomical Region | Hemi<br>(L/R) | Cluster<br>Size<br>(voxels) | *p-value* | *t-value* | MNI Coordinates of Cluster Peak |  |  |
| --- | --- | --- | --- | --- | --- | --- | --- |
|  |  |  |  |  | X | Y | Z |
| **Lingual Gyrus\*** | **R** | **217** | **0.006** | **5.62** | **12** | **-66** | **-6** |
| Intracalcarine Cortex | R |  |  |  |  |  |  |
| Occipital Fusiform Gyrus | R |  |  |  |  |  |  |
| Temporal Occipital Fusiform Cortex | R |  |  |  |  |  |  |
| Precuneous Cortex | R |  |  |  |  |  |  |
| **Cuneal Cortex\*** | **L** | **1446** | **0.002** | **5.11** | **-8** | **-82** | **20** |
| Cuneal Cortex | R |  |  |  |  |  |  |
| Lateral Occipital Cortex sup div (inc. anterior intraparietal sulcus hiP1&3<sup>a</sup>) | L |  |  |  |  |  |  |
| Precuneous Cortex | L & R |  |  |  |  |  |  |
| Occipital Pole | L & R |  |  |  |  |  |  |
| Supracalcarine Cortex | L & R |  |  |  |  |  |  |
| Intrcalcarine Cortex | L |  |  |  |  |  |  |
| Superior Parietal Lobule (inc. anterior intraparietal sulcus hiP1&3<sup>a</sup>) | L |  |  |  |  |  |  |
| Angular Gyrus | L |  |  |  |  |  |  |
| Lingual Gyrus | L |  |  |  |  |  |  |
| **Lateral Occipital Cortex\* (inc. anterior intraparietal sulcus hiP1&3<sup>a</sup>)** | **L** | **76** | **0.017** | **4.91** | **-42** | **-64** | **34** |
| Angular Gyrus | L |  |  |  |  |  |  |
| Supramarginal Gyrus, posterior division | L |  |  |  |  |  |  |

*Note.* Anatomical labels taken from the Harvard-Oxford (HO) Atlas bundled with FSL 6.0.0. \*Cluster peak. <sup>a</sup>Reflects anatomical label from Jeulich Histological Atlas available with FSL 6.0.0. p- and t-values reflect cluster peak. Labels are only reported from regions which survived correction at the voxel level, and subsequent Bonferroni correction.

To understand the function of these structural correlates of MA, a further GLM was constructed comprising the Math-AS scores and nuisance variables as outlined previously, plus those variables that had earlier shown the most robust associations with the level of MA: attentional control (SHIFT), online memory (WORKING MEMORY) and mathematical achievement (WJ). This GLM was applied to those voxels that were previously identified as being negatively associated with MA.

When controlling for attention, working memory and mathematical achievement, a large number of those voxels initially found to be negatively associated with MA were no longer statistically significant, particularly within the IPS. Although this result alone is insufficient to determine a difference, the data suggest that the association between MA and GMV in these voxels may be mediated by one or more of the newly modelled variables. To assess this possibility, the linear directional contrasts for each newly modelled variable was computed. These data suggested that, of the additional variables added, attentional control was negatively associated with a large proportion of those voxels that were no longer associated with MA (see *Supplementary Information: Fig. S2*). However, this result did not survive correction for multiple-comparisons. Working memory and math achievement did not explain the association with MA (see *Supplementary Information for detailed modelling procedure*).

To further assess this, the mean GMV was extracted for each of the 3 clusters identified in the earlier VBM analysis. Though large, the spatial arrangement of these clusters may demarcate differentiation of function, as each comprised relatively distinct anatomical regions: lingual gyrus, cuneal cortex and the intraparietal sulcus. SHIFT, WORKING MEMORY and WJ were entered into a series of partial correlations to examine their relationship with GMV within each of the 3 clusters. When controlling for age, biological sex and Math-AS score, attentional control (SHIFT) was shown to be negatively associated with GMV across the IPS, although this result did not survive a Bonferroni correction for 9 tests. All other tests were non-significant at the uncorrected level (see Table 4).

**Table 4.**
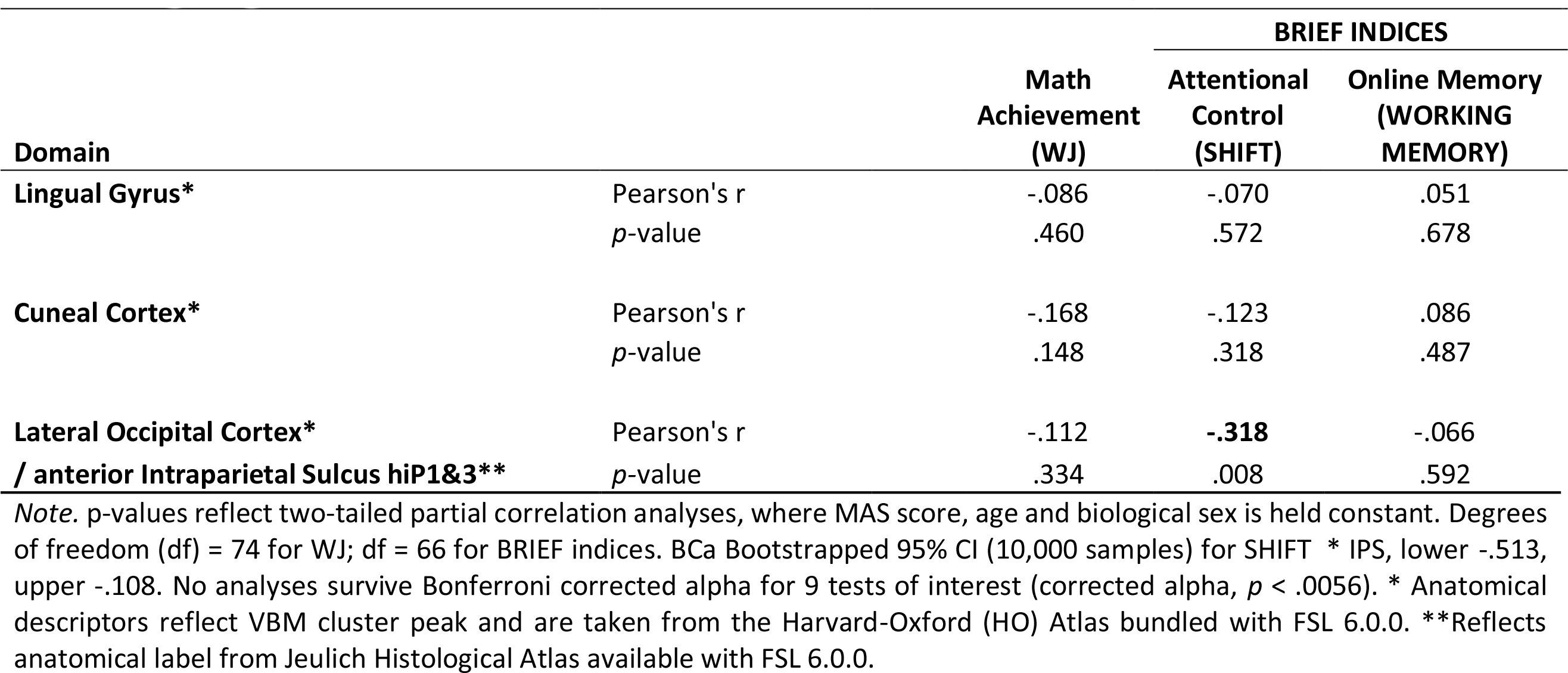
Partial Correlation Results for Regional Grey Matter Volume, Numerical and Cognitive Indices

### The Cognitive Architecture of MA and Resultant Outcomes in Numerical Achievement

Suárez-Pellicioni and colleagues^10^ outline a model in which intrusive thoughts resultant from MA consume working memory. This in turn, is argued, to expend already limited attentional resources in high MA individuals, leading to diminished performance when complex mathematical operations are performed. Such a relationship provides a causal explanation for the association between MA (Math-AS) and mathematical achievement (WJ). We tested this theoretical model using mediation analysis. When controlling for working memory and the nuisance variables age and biological sex, the relationship between MA and math achievement was no longer significant (c’) (see Table 5 and Fig. 2). The overall mediation model found that MA, working memory, age and biological sex explained approximately 55% of the variance in math achievement, (R^2^ =.5527, F(4, 66) = 20.39, p <.0001). The mediation model suggested that higher levels of MA (Math-AS) were associated with slightly elevated difficulty with holding appropriate information in mind (WORKING MEMORY), which in turn resulted in reduced math achievement (WJ).

**Figure 2.**
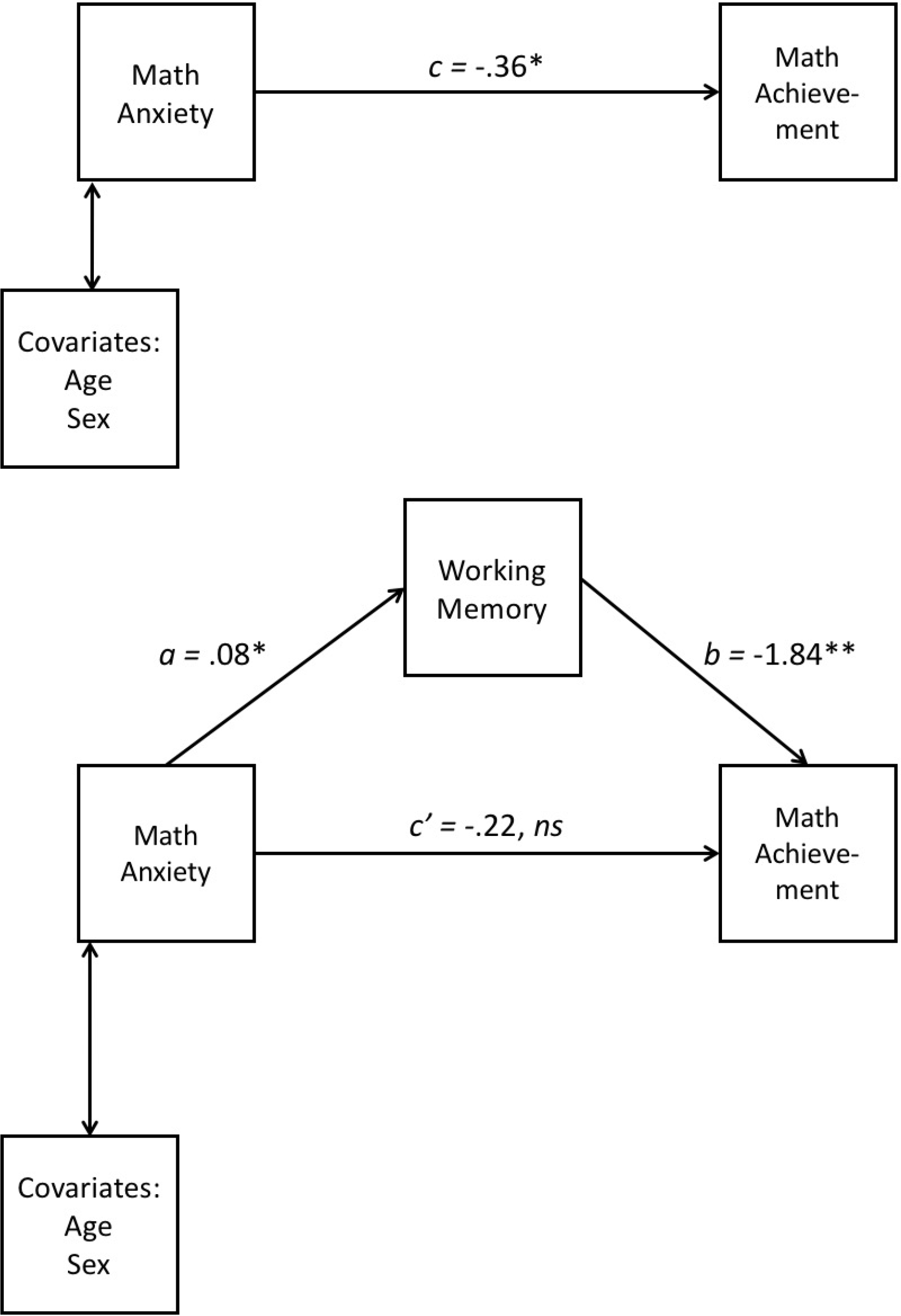
Summary coefficients for mediation model. *Note.* Path a = unstandardised IV to mediator; path b = unstandardised mediator to DV; path c = unstandardised total effect (IV to DV); path c’ = unstandardised direct effect. Coefficient values rounded to 2 decimal places; full values reported in accompanying table (3). ns = non-significant * p< 0.01; ** p<0.001.

**Table 5.**
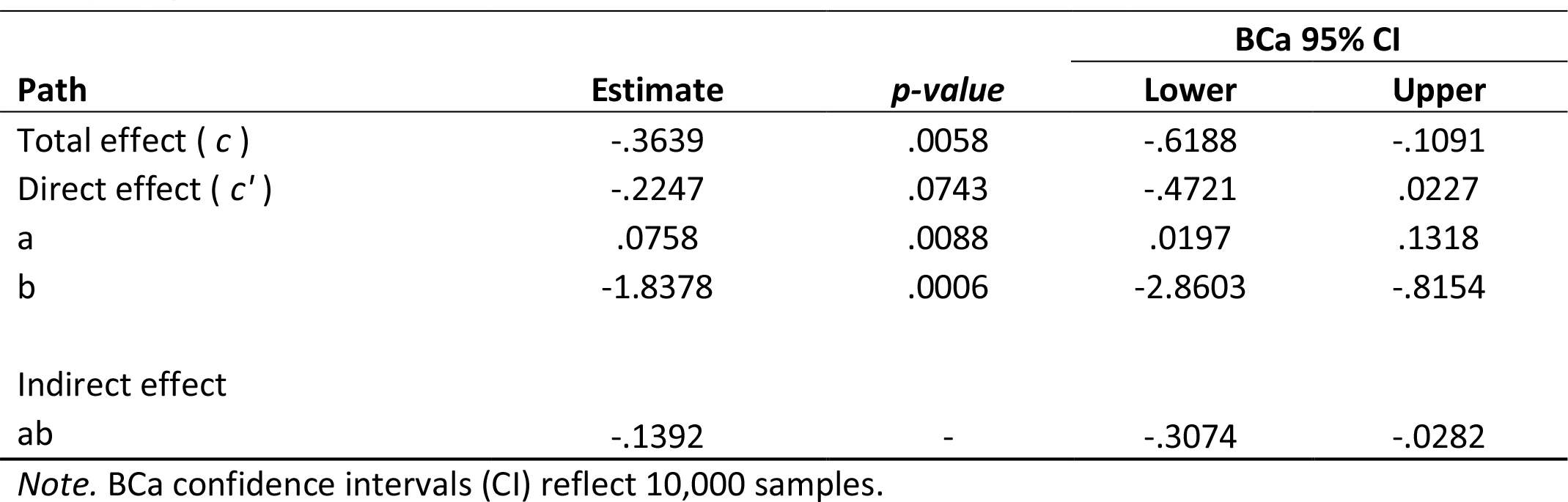
Mediation Path Coefficients and Confidence Intervals for Math Anxiety Predicting Math Achievement

## Discussion

The current study sought to examine the neurocognitive bases of MA in typically developing children. Unlike most prior work, however, the analyses focused on the level of MA, which is a more rigorous approach than making comparisons based on the presence or absence or of it^16^. This work therefore provides both a description of how MA might manifest itself across a cohort of typical children, as well as a detailed evaluation of theoretically driven factors that might influence the degree of MA.

Most of the children in the sample demonstrated low-levels of MA. Around 10% of the entire sample reported moderate to high-levels of MA and prevalence increased linearly with age. Aside from numerical representations, all of the other cognitive variables that were of theoretical interest were associated with the degree of MA: the more math anxious children demonstrated reduced inhibitory and attentional control, as well as lower working memory capacity and math achievement. Though MA appears to be associated with differences in baseline executive functions in children with MA, working memory was also shown to mediate the relationship between MA and math achievement suggesting that, possibly in addition to baseline differences, capacity issues are exacerbated ‘online’ when working with mathematics by the physiological response to anxiety, which in turn leads to poorer learning and performance in mathematics.

Contrary to previous studies^17,18^, we found no association between MA and low-level numerical representations. There has been considerable debate regarding the presence of a low-level deficit in math ability in MA^10^. It may be that the task used here was not fine-grained enough to highlight any association, although note that the data from the task used did correlate with the measure of math achievement, suggesting that the task was tapping into numerical cognition to some degree. The current pattern of results is also indicative of experiential differences between high and low math anxious individuals, as MA may result in avoidance of exposure to math problems^19^. Moreover, an experientially-driven reduction in math achievement does not discount a primary, causal deficit in numerical representation; it may be that more extreme levels of MA reflect both causal (representational) and affectual (experiential) mechanisms. Indeed, cross-sectional work strongly suggests the presence of a less precise representation of numerical representation in adults with high MA.^18^

The present study also identified that reduced grey matter volume in the IPS, lingual gyrus and cuneal cortex was associated with increased MA. The identification of a structural correlate in relatively young children suggests that there may be differences in early brain structure that underpin the development of MA. Thus, whilst prior functional imaging studies demonstrate how network functionality can explain differences in mathematical performance in people with math anxiety^8,9^, these data provide the first evidence of a possible underlying structural basis. Genetic modelling suggests that around 40% of variance in MA^5^ can be explained by heredity, thus, a neurodevelopmental precursor is not implausible. Prior work has implicated the IPS in a network of regions showing aberrant activity in young, math anxious children, where these patterns of activation were unrelated to intelligence, general anxiety, reading ability or working memory.^9^ Our data suggest that the IPS region identified in the current study might serve an attentional function, where children with reduced grey matter in this area had lower reported attentional resources, and this reduced attentional capacity was associated with increased MA. A potential model that should be examined in future research is that children with MA, or who go on to develop MA, start off with differences in IPS structure, which translate into a deficit in baseline attentional capacity. According to this view a limited ability to attend to stimuli, particularly mathematical stimuli – where demands on attentional resources are often high due to the nature of arithmetic problems – could result in general feelings of anxiety, which later become habitually associated with *doing* math, causing the development of MA.

In addition to providing a rich description of the neurocognitive bases of the level of math anxiety, the current study provides testable hypotheses regarding the emergence and maintenance of MA. Longitudinal work is required, however, to assess the validity of these assertions.

## Acknowledgements

Authors CEH FS, and RCK were supported by the European Research Council (Learning and Achievement, award number 338065). Authors LJF and CGS were supported by grant PSI 2014-53427-P from Ministry of Economy and Competitiveness (FEDER funding) and grant 19267/PI/14 from Fundación Séneca.

## Author Contributions Statement

LJF conceived and designed the over-arching study. All authors contributed to the math anxiety study concept. FHS translated the Math-AS into Spanish and organized and interpreted the Math-AS scores. CGS organized and interpreted the BRIEF questionnaire. LJF, AI and JMGS designed and implemented the MRI protocol. LJF, AI, FHS, CGS and JMGS oversaw the testing and data collection. CEH, CYL, FS and RCK were given access to the subset of data described in this article to examine math anxiety, following LJF taking a sabbatical within RCK’s laboratory at the Department of Experimental Psychology, University of Oxford. CYL contributed to the organization of test measures and raw data. CEH, CYL and FS conducted the behavioural analysis. CEH conducted the neuroimaging analyses. CEH and CYL drafted the manuscript. LJF, RCK, FS, AI, FHS, CGS and JMGS provided critical revisions of the manuscript. All authors approved the final version of the manuscript for submission.

## Additional Information

### Data Availability

The research data supporting this publication are available on the Open Science Framework repository, see DOI 10.17605/OSF.IO/PDFJE. The senior author, LJF, may be contacted regarding the wider dataset.

